# Permeabilization with fenchone enhances cryopreservation of *Drosophila* embryos

**DOI:** 10.1101/2025.11.06.687078

**Authors:** David M. Zimmerman, Rachel Yin, Madyson Vaca, Aravinthan D. T. Samuel, Benjamin L. de Bivort

## Abstract

The difficulty of cryopreservation has long been a limitation of *Drosophila melanogaster* as a genetic model organism. Here we report a statistically significant improvement in the efficiency of *Drosophila* cryopreservation by substituting limonene with the monoterpenoid fenchone in the embryo permeabilization step of a previously published method. We found that fenchone-permeabilized embryos exhibit greater uptake of cryoprotectant compared with those permeabilized by limonene, and a ~6-fold increase in the rate of egg-to-adult survival for wild-type flies. Using this improved protocol, we successfully cryopreserved and revived precious strains after 12 months of storage in liquid nitrogen. These results suggest that fenchone is a superior permeabilizing agent for fly embryo cryopreservation, expanding possibilities for the long-term maintenance of *Drosophila* and other insect species. Further refinement of this approach may enable cryopreservation to replace continuous culture as the method of choice for routine maintenance of fly stocks.

## Introduction

The fruit fly *Drosophila melanogaster* is an important model organism to many areas of biology. The fly’s compact size, short generation time, extensive genetic toolkit, and general hardiness have made it a powerful experimental platform for studies of development, ecology and evolution, physiology, and behavior. Unfortunately, unlike other widely used model organisms such as *E. coli* (Sprouffske et al., 2016), budding yeast (Bond, 2007), *C. elegans* (Barranco and Risco, 2022; O’Connell, 2022), and even some vertebrates (Mandawala et al., 2016), the fruit fly has long proven recalcitrant to cryopreservation.

Simple freezing of embryos, larvae, and adults normally leads to complete lethality upon thawing, with the result that continuous culture (“flipping”) remains the only practical method of maintaining *Drosophila* strains in the laboratory. Indeed the amount of labor involved in flipping hundreds or thousands of different strains on a regular basis imposes a considerable burden on even well-resourced fly labs. In addition, continuous culture inevitably leads to the accumulation of genetic mutations over time, which can present significant problems for rigorous studies of population and behavioral genetics (Burke and Rose, 2009; Houot et al., 2010; Colomb and Brembs, 2014; Sprouffske et al., 2016). Beyond *Drosophila*, there is growing interest in developing reliable cryopreservation methods for non-model insects, to facilitate genetic and ecological studies of agriculturally and medically relevant species.

Despite reports of successful cryopreservation of *Drosophila* embryos (Steponkus et al., 1990; Mazur et al., 1992) and germ cells (Nishimura et al., 2023), no technique has achieved sufficient ease of use and effectiveness to earn widespread adoption. Recently, Zhan et al. (2021) reported a new vitrification-based cryopreservation method with egg-to-adult survival rates ranging from 10 to 75%. This approach involves loading embryos with the cryoprotectant ethylene glycol via treatment with the monoterpene solvent D-limonene, which has previously been shown to function as an effective permeabilizing agent for *Drosophila* embryos (Rand et al., 2010). Similar vitrification-based strategies have also been used successfully for cryopreservation of various non-model Diptera (Rajamohan et al., 2003; Rajamohan and Leopold, 2007; Augustinos et al., 2016). Such an efficient method has the potential to revolutionize the laboratory maintenance of *Drosophila* strains and obviate one of the organism’s greatest weaknesses as an experimental model system. Our attempts to implement this method did achieve successful cryopreservation, but with much lower egg-to-adult survival. Our best results using limonene as the permeabilizing agent yielded survival rates in the range of 0.5–2%. However, we serendipitously discovered that substituting limonene with the alternative monoterpenoid fenchone resulted in survival rates as high as 12 ± 2%, representing an order-of-magnitude improvement in wild-type flies. This increase in efficiency could be of practical significance to other *Drosophila* workers as well as biologists concerned with the cryopreservation of similarly recalcitrant insect species.

## Materials and methods

### Fly husbandry

Flies were raised in 6 oz. bottles of standard cornmeal-based “white” food at room temperature (22 °C). All experiments comparing the efficacy of limonene and fenchone were conducted with flies of the de Bivort lab’s wild-type Canton-S stock. Long-term cryopreservation experiments used outbred flies from two synthetic populations (gift of S. Lall). These populations were constructed from 40 DGRP lines (Mackay et al., 2012) that were (1) mated in round-robin fashion for four generations, (2) maintained in large cage populations for 12 generations, and (3) subjected to artificial selection for behavioral variability for 22 generations. As such, they represent precious flies of an unstable genetic composition: ideal candidates for cryopreservation.

### Embryo collection

To collect embryos for cryopreservation, 600–1000 flies of mixed sex were collected 5 to 8 days after eclosion and allowed to lay eggs for 1 hour at 25 °C on grape juice agar plates supplemented with fresh yeast paste. The same adult flies were used for up to 5 consecutive collections on fresh grape plates. Eggs from the first 1-hour collection were discarded, as this initial laying period serves to clear retained eggs from females and yields more developmentally synchronized embryos in subsequent collections. Collections containing fewer than approximately 200 eggs were discarded. Each collection was treated as an independent replicate. Where possible, different collections from the same batch of parental flies were assigned to different treatment groups (e.g., fenchone vs. limonene) to minimize confounding effects of parental batch on treatment outcome. Eggs were allowed to develop for 22 hours at 20 °C, until a plurality of the population is at stage 16.

### Cryopreservation and storage

We followed the detailed cryopreservation protocol of Zhan et al. (2021) except for modifications to the permeabilization step (Fig. 1). In brief, embryos were dechorionated with bleach; permeabilized by successive incubations in isopropanol, limonene or fenchone (dissolved in heptane), and pure heptane; dispersed into a floating monolayer using a paintbrush; cryoprotected by successive incubations in 13% (w/v) ethylene glycol and 39% (w/v) ethylene glycol + 9% (w/v) sorbitol; transferred to a nylon mesh substrate; and plunge-frozen in liquid nitrogen (LN2). Permeabilization was performed either with the original D-limonene/heptane (1:4 [v/v]) mixture or with an analogous mixture of D-fenchone/heptane (1:4 [v/v]). The length of the incubation in this solution was varied systematically from 10 s to 30 s. D-limonene (CAS 5989-27-5, product no. 183164), D-fenchone (CAS 4695-62-9, product no. 46210), and heptane (CAS 142-82-5, product no. 34873) were from Millipore Sigma.

**Figure 1.**
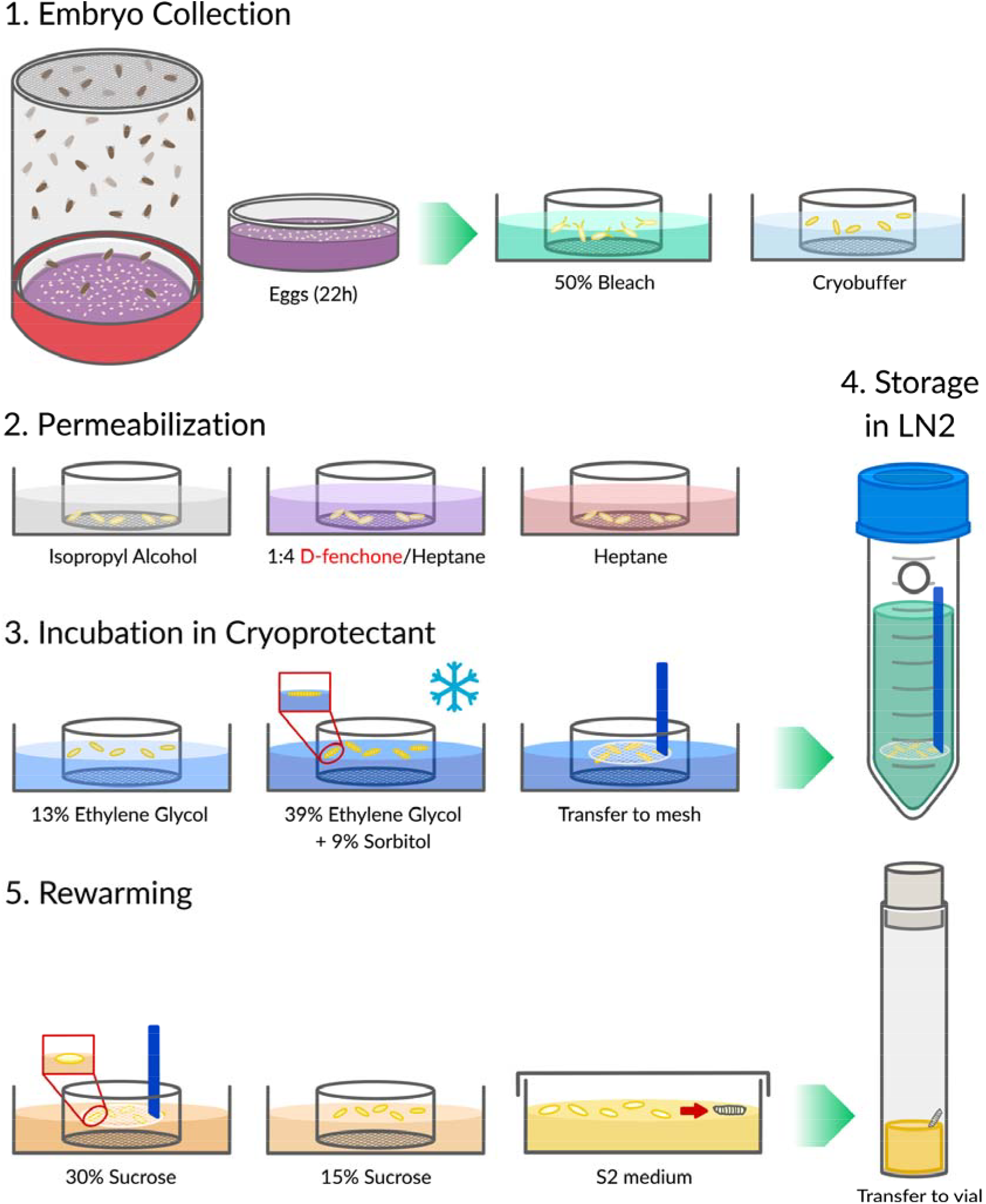
Overview of cryopreservation technique. Schematic overview of vitrification-based cryopreservation technique. Eggs are collected on grape juice plates and dechorionated in bleach; the dechorionated embryos are transferred to cryobuffer. Embryos are then permeabilized (by D-fenchone, rather than D-limonene, dissolved in heptane); loaded with ethylene glycol cryoprotectant; and vitrified by plunge-freezing in LN2 on a nylon mesh substrate. Embryos are maintained in LN2 for long-term storage, prior to rewarming in sucrose, overnight recovery in S2 medium, and transfer to cornmeal food. Except for the substitution of fenchone for limonene, the procedure is substantially the same as in Zhan et al. (2021).

After immersion in LN2, cryomeshes were transferred without exposure to air into 50 mL conical tubes for long-term storage. Prior to use, holes were made in the sides of these tubes with a hot metal rod to prevent explosive buildup of nitrogen vapor. Conical tubes were kept in LN2 inside a closed storage dewar prior to rewarming in sucrose, following the procedure of Zhan et al. (2021). Rewarmed embryos were kept in sterile S2 medium at 22 °C until hatching, then transferred as larvae to cornmeal food in vials. Surviving animals were counted manually at the appropriate developmental stage.

### Dye uptake assay

The same procedure as described by Zhan et al. (2021) was used to assess the efficiency of permeabilization. Briefly, permeabilized embryos were incubated for 5 min in 0.1% rhodamine dye (CAS 81-88-9, Millipore Sigma, product no. R6626) dissolved in cryobuffer, followed by visualization on an upright epifluorescence microscope equipped with a 1X/0.04 air objective and a standard RFP/TRITC filter set. Eggs from 3 independent collections were mixed before being randomly distributed between the three experimental groups. Images were analyzed with Cellpose and scikit-image to segment individual embryos and compute the average fluorescence intensity of each embryo (van der Walt et al., 2014; Stringer et al., 2021).

### Statistical analysis

Survival data were modeled with logistic regression. In particular, the larval hatch rate data in Fig. 2a were fit to a binomial generalized linear mixed model with logit link, using the lme4 R package for maximum likelihood estimation; a random-intercept term (normally distributed on the logit scale) captured batch-related overdispersion. As the adult survival data in Fig. 2b exhibited minimal overdispersion, the random effect was omitted. However, to deal with the problem of separation caused by the zero counts in the vehicle (i.e., pure heptane, with neither limonene nor fenchone) treatment group, the adult GLM was fitted using Firth’s bias reduction method as implemented by the brglm package. In both cases, t-tests were used for pairwise comparisons of the estimated regression coefficients. In Fig. 2e, pairwise differences (in the sense of stochastic dominance) with respect to the typical mean fluorescence intensity were evaluated using Brunner–Munzel tests.

**Figure 2.**
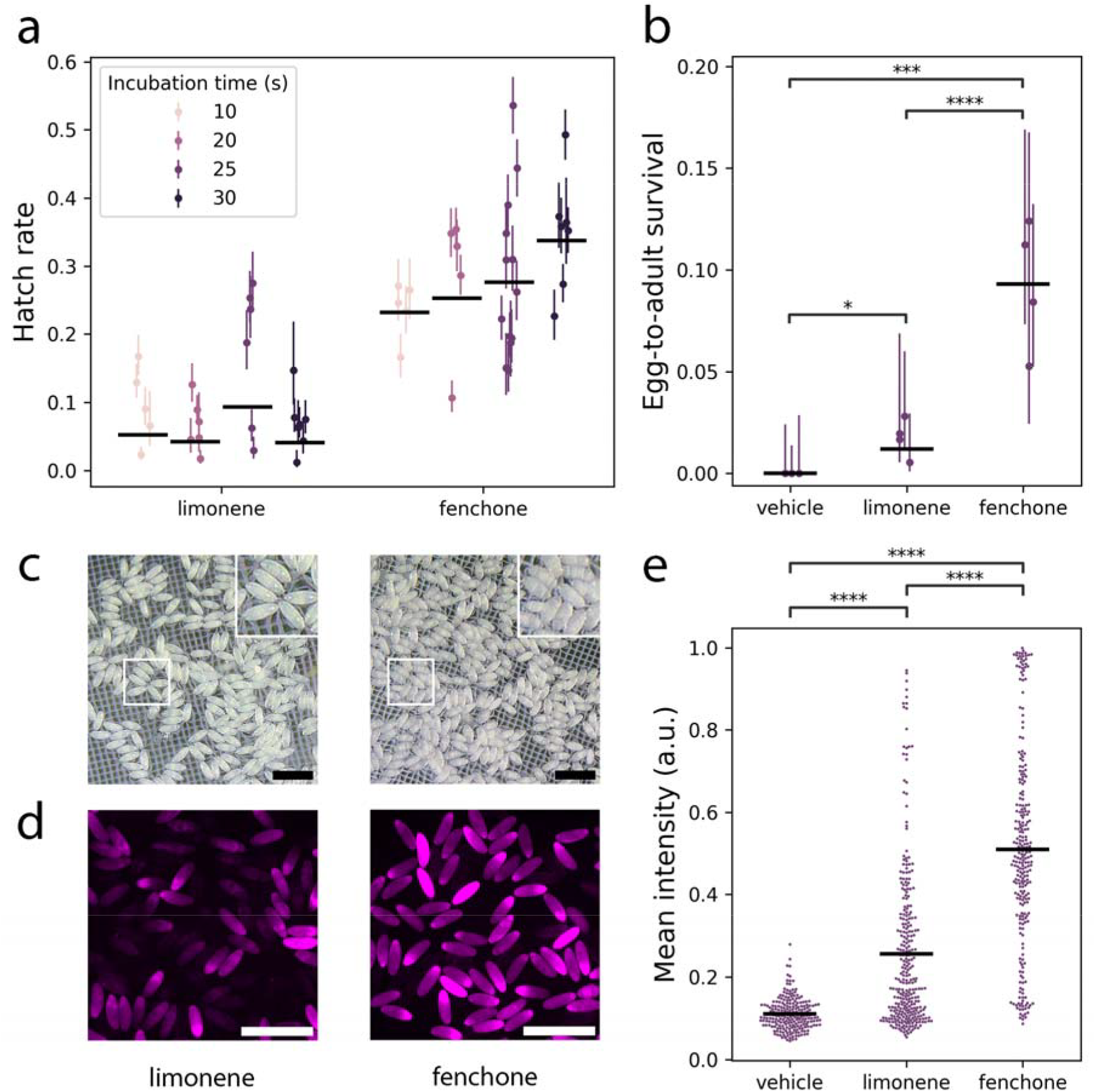
Fenchone achieves superior permeabilization and post-freezing survival. (a) Egg-to-larva hatch rate vs. incubation time for batches of embryos permeabilized with either limonene or fenchone (in heptane). The marginal mean hatch rate is significantly greater for fenchone than for limonene (*p* < 0.0001). Horizontal lines represent the mean hatch rate for each condition, weighted by the binomial precision of individual measurements. (b)Egg-to-adult survival rate for embryos permeabilized with limonene, fenchone, or vehicle (heptane) for 25 s. Horizontal lines represent the mean hatch rate for each condition, weighted by the binomial precision of individual measurements. Asterisks indicate significant mean differences: *, *p* < 0.05; ***, *p* < 0.001; ****, *p* < 0.0001. (c)Reflected-light images of embryos showing shrinkage after incubation in either limonene (left) or fenchone (right) for 3 min. Scale = 1 mm. Inset shows detail of embryo shrinkage, magnified 1.8X. (d)Epifluorescence images of embryos showing differential uptake of rhodamine dye following permeabilization with either limonene (left) or fenchone (right) for 25 s. Scale = 1 mm. (e)Mean fluorescence intensity of embryos permeabilized with limonene, fenchone, or vehicle (heptane) for 25 s and incubated in rhodamine dye for 5 min. Horizontal lines represent the overall mean for each condition. Asterisks (****) indicate significant pairwise differences on the Brunner– Munzel test, *p* < 0.0001.

## Results and discussion

### Fenchone promotes embryo permeabilization and survival following cryopreservation

Motivated by a serendipitous accident in which replacing limonene with fenchone led to qualitatively enhanced survival following cryopreservation, we set out to replicate this result and potentially optimize a fenchone-based cryopreservation protocol (Fig. 1). We tested the effect of different periods of incubation in 1:4 limonene/heptane vs. 1:4 fenchone/heptane solution on post-freezing embryo-to-larva hatch rate (Fig. 2a). Embryos permeabilized with fenchone survived to the larval stage at significantly higher rates than did embryos permeabilized with limonene (0.28 ± 0.02 vs. 0.08 ± 0.009 [mean ± s.e.m.], *p* < 0.0001). Fenchone’s effect on post-freezing hatch rate varied more with incubation time than did limonene’s effect (χ^2^(3) = 9.19, *p* = 0.027), exhibiting a non-significant trend toward higher effectiveness at longer incubation periods. In contrast, limonene produced a higher hatch rate at an incubation period of 25 s than at either 20 s (*p*_adj_ = 0.026, Bonferroni-corrected) or 30 s (*p*_adj_ = 0.028). We next compared the effect of permeabilization with limonene vs. fenchone on egg-to-adult survival, using a fixed incubation period of 25 s for both reagents. Consistent with its observed effect on larval hatch rates, fenchone also promoted significantly increased post-freezing egg-to-adult survival relative to limonene (0.101 ± 0.011 vs. 0.018 ± 0.005, p < 0.0001; Fig. 2b). From these experiments, we conclude that fenchone is superior to limonene as a permeabilizing agent for cryopreservation.

Successfully permeabilized embryos shrink due to osmosis when incubated in a 13% solution of the cryoprotectant ethylene glycol (Zhan et al., 2021). We found that permeabilization with fenchone results in more robust shrinkage of embryos during this step (Fig. 2c). This observation suggested to us that the effect of fenchone on survival rate might be mediated by an increase in the efficiency of permeabilization, leading in turn to enhanced uptake of the cryoprotectant agent. Consistent with this hypothesis, we determined that fenchone-permeabilized embryos fluoresce more brightly on average after a 5 min incubation in rhodamine dye than do limonene-permeabilized embryos (Fig. 2d–e; *p* < 0.0001).

Finally, as a concrete demonstration of the types of experiments made possible by the use of fenchone for permeabilization, we cryopreserved two precious *Drosophila* strains that were products of a 21-generation artificial selection experiment. We were able to revive both of these strains after storing them for 12 months in LN2. Crucially, the survival rates on rewarming (Table 1) were comparable to those we achieved for the Canton-S flies used in the experiments described above, which were rewarmed immediately after freezing.

**Table 1.** Egg-to-adult survival for lines VC3 and VS3, thawed after 1 year in LN2 storage (Sep 2023 to Sep 2024).

| Strain | Batch | Total eggs | Survivors | Survival rate |
| --- | --- | --- | --- | --- |
| VC3 | 1 | 137 | 17 | 0.124 |
| VC3 | 2 | 180 | 25 | 0.139 |
| VC3 | 3 | 182 | 15 | 0.082 |
| VS3 | 1 | 329 | 140 | 0.426 |

### Mechanistic considerations and prospects for further improvement

The biophysical mechanism by which terpenes and terpenoids such as limonene and fenchone permeabilize cellsis incompletely understood (Abramov et al., 2001; Espina et al., 2013; Ortiz et al., 2024). It has been suggested that permeability of *Drosophila* embryos is mainly limited by the thin waxy layer (~5 nm thick) that encases the vitelline membrane (Rand et al., 2010). Limonene, which is used in various household cleaners as a non-toxic solvent for the removal of waxy substances, was first employed as an embryo permeabilizing agent on the hypothesis that it might offer a less toxic and water-miscible alternative to linear hydrocarbons such as heptane and octane. The physicochemical properties of fenchone (e.g., its polar ketone group) may improve upon the effectiveness of limonene for solubilizing the waxy layer while presenting a similarly low level of toxicity. Nevertheless, compounds in this family are known to exert at least some toxic effects on the normal growth and development of *Drosophila* spp. when administered for long enough periods at high enough doses (de Souza et al., 2024). It is likely that a more systematic comparison of different compounds in this family would reveal agents with even better properties for cryopreservation applications. That at least one such compound, D-fenchone, leads to a significant increase in the efficiency of *Drosophila* cryopreservation suggests that further improvement is possible.

At present, we feel that cryopreservation is already sufficiently effective for use with precious and/or difficult-to-maintain strains, for which the intrinsic risks of loss or genetic drift associated with live culture justify the extra effort of this technique. However, cryopreservation is probably still too difficult and time-consuming to replace flipping for most applications. Nevertheless, we are optimistic that further optimization will make cryopreservation a viable replacement for routine stock maintenance, akin to how bacteria and nematodes are routinely frozen for long-term storage. In addition, improvements in embryo permeabilization could facilitate the application of vitrification-based cryopreservation to a wider variety of non-model insects, where stable preservation of wild or genetically unique populations would be of great value for both research and conservation.

## Data accessibility

Data and statistical analysis code are available from Zenodo at https://doi.org/10.5281/zenodo.17543286 (Zimmerman et al., 2025).

